# Rapid sensory integration in orexin neurons governs probability of future movements

**DOI:** 10.1101/620096

**Authors:** Mahesh M. Karnani, Cornelia Schöne, Edward F. Bracey, J. Antonio González, Paulius Viskaitis, Antoine Adamantidis, Denis Burdakov

**Affiliations:** Institute for Neuroscience, Department of Health Sciences and Technology, ETHZ, Switzerland; The Francis Crick Institute, London, UK; Institute of Psychiatry, Psychology & Neuroscience, King’s College London, UK; Department of Neurology, Inselspital, University of Bern, Switzerland; The Rowett Institute, School of Medicine, Medical Sciences and Nutrition, University of Aberdeen, UK

## Abstract

Internally and externally triggered movement is crucial for survival and is controlled by multiple interconnected neuronal populations spanning many brain areas. Lateral hypothalamic orexin/hypocretin neurons are thought to play a slow, modulatory part in this scheme through their key role in promoting metabolism and wakefulness. However, it is unknown whether orexin/hypocretin neurons participate in rapid neural processing, leading to immediate movement. Furthermore, their role in sensorimotor transformations remains unknown. Here we use cellular-resolution Ca^2+^ imaging and optogenetics to show that orexin/hypocretin cells are instantaneous regulators of self-generated and sensory-evoked movement. They are activated before and during movement, preventing this activation reduces self-generated locomotion, and optogenetic mimicry of this transient activation rapidly initiates locomotion. We find that the same orexin neurons whose activity predicts movement initiation are rapidly controllable by external sensory stimuli, and silencing orexin cells during sensation prevents normal motor performance. These findings place orexin neurons in a physiological position of unexpectedly rapid and strong sensorimotor control.

## Introduction

A core function of the nervous system is to detect and identify salient sensory inputs and generate motor signals that link sensation to action. The importance of the neocortex, and other classical sensory and motor brain regions, in these rapid sensorimotor transformations is well established (Crochet et al., 2019; Svoboda and Li, 2018). While neurons in the lateral hypothalamus (LH) also show fast sensory responses (Burton et al., 1976; Mora et al., 1976; Rolls et al., 1976), and are known to promote movement (Kosse et al., 2017; Li et al., 2018; Sinnamon, 1993; Yamanaka et al., 2003), their role in rapid sensorimotor processing is mostly unstudied. Instead, a large body of literature on the hypothalamus has focused mainly on control of hormone secretion, slow neuropeptide output, and long-lasting states like sleep or obesity, measured at the resolution of hours, and almost always using chronic manipulations of hypothalamic neuronal activity. This has reinforced a prevailing dogma that the primary function of the LH is slow regulation of feeding, arousal and metabolism (Arrigoni et al., 2018; Saper et al., 2005).

A key genetically-distinct population of LH neurons that responds to external stimuli are the orexin (hypocretin) neurons, which are evolutionarily conserved and project throughout the brain (Tyree et al., 2018). Orexin neurons are critical regulators of arousal and whole body metabolism (Adamantidis et al., 2007; Yamanaka et al., 2003), that are commonly thought to act on a slow timescale. However, unit recordings (Hassani et al., 2016; Lee et al., 2005; Mileykovskiy et al., 2005; Takahashi et al., 2008) and averaged population recordings with fiber photometry (González et al., 2016a; Inutsuka et al., 2016) have shown that orexin neurons respond to sensory stimuli, and that they can change their spike output on the millisecond timescale. Because the activity of orexin cell ensembles has not been recorded before at single-cell spatial resolution, it is unclear what proportion of orexin neurons are responsive and how this activity is coordinated between them. Furthermore, the physiological significance of their responses is unclear. We therefore set out to assess the behavioural correlates of orexin neuron activity dynamics, and how rapidly orexin neurons can control behaviour in response to sensory input.

We used two-photon volumetric imaging to record rapid dynamics across hundreds of orexin neurons in behaving mice. This revealed numerous aspects of their instantaneous activity modulation including a new functional subtype division within the population. We observed that individual orexin cells are activated within tens of milliseconds after sensory stimuli. Many orexin cells are also activated seconds before self-paced movement initiation (self-initiated locomotion). Loss- and gain-of-function manipulations revealed that activating orexin cells leads to locomotion within hundreds of milliseconds, and inhibiting orexin cell activity decreases the likelihood of self-initiated locomotion. Finally, temporally-precise optogenetic silencing, we provide causal evidence for the role of orexin cells in rapid neural transformations from sensory input to movement output.

## Results

Orexin neurons are often studied in a slow framework where emphasis is placed on modulation of long-lasting states or regulation of ‘background’ homeostatic variables. If orexin neurons only regulate output variables slowly, they would not need to undergo rapid activity changes on a subsecond timescale and manipulating their activity should not affect processes on the subsecond timescale. In order to probe how fast their activity is modulated, we set out to record orexin neurons in awake behaving mice. We recorded the activity of orexin neurons through the fluorescence of the GCaMP6s Ca^2+^ sensor (Chen et al., 2013) which was delivered in a vector under the orexin promoter, restricting expression to orexin cells (cell counts in Methods)(González et al., 2016b). Simultaneous electrical and optical recordings in brain slices showed that GCaMP6s fluorescence faithfully and rapidly reported spiking activity of orexin cells (Figure 1A-E). Using implanted graded refractive index (GRIN, Supplementary figure 1) lenses and a miniature fluorescence microscope mounted on the skull, we then recorded activity of individual orexin neurons in freely behaving mice as they navigated a familiar 24×24cm arena that contained a food pod and an opposite sex conspecific (Figure 1F-J). It was immediately evident that orexin neurons do undergo rapid activity modulation time-locked to natural behaviours lasting only a few seconds (Figure 1J).

**Figure 1.**
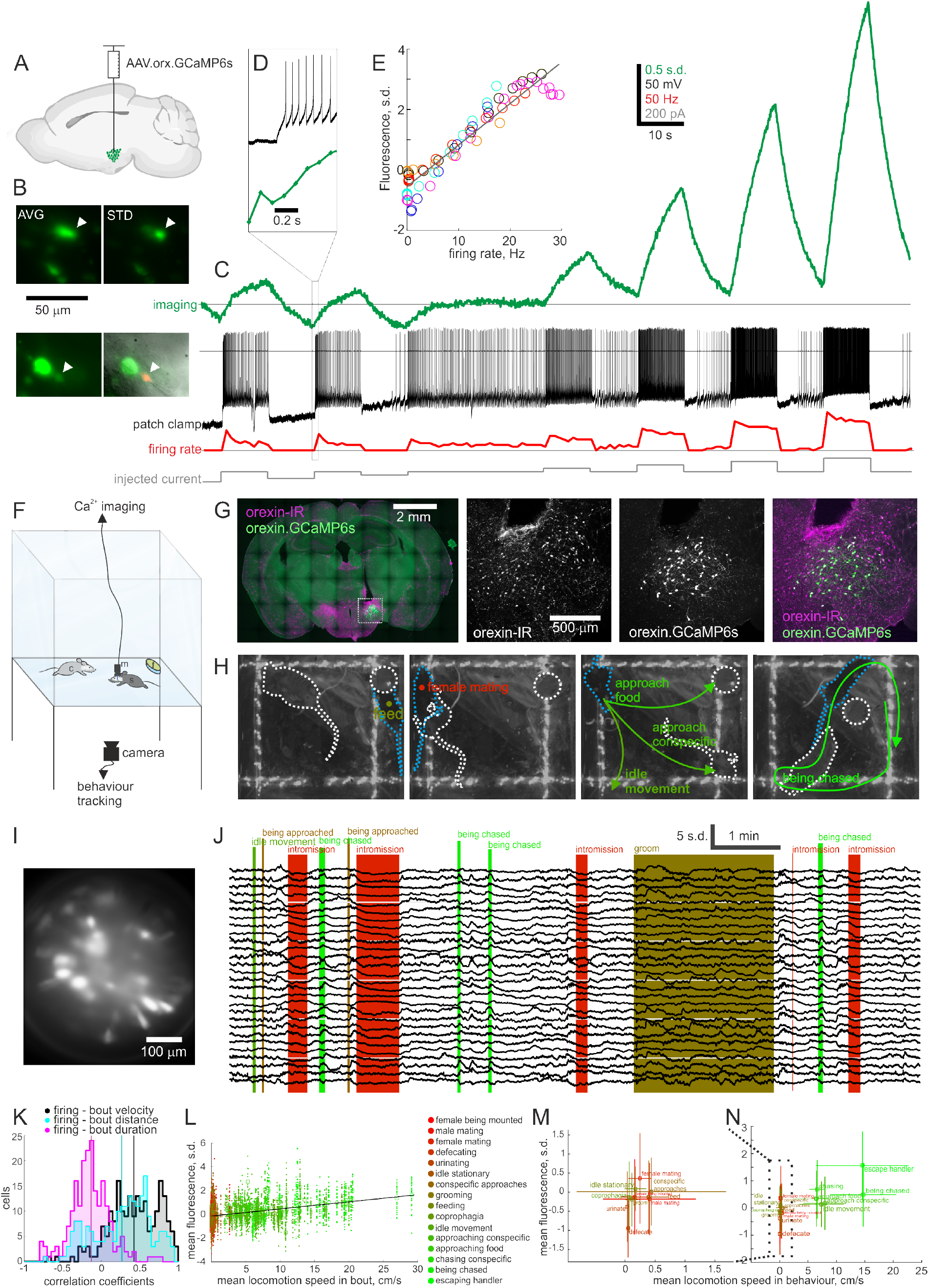
Orexin neuron Ca imaging in freely moving socializing animals. A, A virus was used to drive GCaMP6s expression under the orexin promoter, such that only orexin cells express GCaMP6s. B, An orexin neuron in a brain slice was imaged with a miniature endoscope at an angle (above micrographs: AVG, average time projection; STD, standard deviation projection; recorded neuron at arrow head) and with patch clamp microscope from above (below left, GCaMP6s fluorescence; below right, merged GCaMP6s, Alexa-568 red patch pipette filling solution and oblique illumination micrograph showing patch pipette; recorded neuron at arrow head). C, Ca-imaging (green, Z-scored ΔF/F_0_), electrophysiology (black) and firing rate (red) example data from cell in B. D, Expanded view of firing burst onset showing short latency from firing burst to Ca increase. E, Combined data across 6 recordings (each recording in a different colour) showing linear relation between maximal fluorescence and firing rate during each 10s current step. R^2^ of linear fit is 0.91. F, Schematic of freely-moving in vivo paradigm where orexin neurons in subject (s) were recorded with miniature endoscope (m) in an arena with a conspecific (c) and food pod (f) while behaviour was tracked from below through the transparent floor. G, Representative micrographs showing orexin immuno-reactivity (IR) and GCaMP6s in coronal sections prepared from an experimental subject. GRIN lens track is visible dorsal to the orexin neurons. H, Representative behaviour video frames showing a female subject (black fur, blue dashed outline) feeding (left), mating (second from left), 3 different medium speed movements (second from right) and being chased (right) by a male conspecific (brown fur, white dashed outline). I, Representative average time projection of Ca-imaging data through the cylindrical GRIN lens. J, Representative time series of z-scored fluorescence (ΔF/F_0_) of 28 orexin neurons across various behavioural epochs. K, Histogram of Pearson correlation coefficients across all recorded cells (mean correlation with bout speed 0.43 ± 0.02; with distance 0.27 ± 0.03; with duration −0.14 ± 0.02; each distribution significantly different from the others, P<10^-17^ with paired Student’s t-test; 207 cells across 12 recording sessions from 5 animals, 2 male, 3 female). L, Mean Ca-activity for each cell during each behaviour bout (points) and their average (squares) coloured by the behaviour class (colour code from red to green follows roughly movement speed in behaviour) plotted as a function of mean locomotion speed in behaviour bout (249 bouts, 5873 cell/bout pairs). Black line is linear fit to bout average data (squares) with R^2^=0.21 and P<10^-13^. M and N, Same data as in L grouped into behaviour classes and plotted as mean ± s.d.

We categorized behaviours into 16 classes that were clearly distinguishable from the camera angle below the animals. The behaviour bouts were rapid, lasting on average 11.2 ± 6.6 s (behaviour, mean duration ± s.d., number of bouts: female being mounted (without intromission, and not leading to it), 3.0 ± 2.0 s, 11; male mating (intromission), 13.8 ± 7.7 s, 15; female mating (intromission), 32.6 ± 29.4 s, 4; defecating, 6.7 ± 1.6 s, 5; urinating, 3.6 ± 1.3 s, 3; idle stationary (no activity or goal detected), 22.5 ± 25.0 s, 19; conspecific approaches, 2.1 ± 2.8 s, 32; grooming, 38.4 ± 39.6 s, 19; feeding, 46.5 ± 38.8 s, 13; coprophagia, 19.4 s, 1; idle movement (self-initiated locomotion without a detected goal), 4.5 ± 2.9 s, 37; approaching conspecific, 3.0 ± 2.0 s, 43; approaching food, 5.2 ± 4.5 s, 22; chasing conspecific, 7.8 ± 7.1 s, 12; being chased (by conspecific), 6.8 ± 4.2 s, 21; escaping handler, 3.6 ± 3.8 s, 6). Across behavioural classes, activity of 33.3% of orexin neurons was strongly correlated (Pearson correlation coefficient >0.6) with the speed of locomotion during the behaviour, 17.4% with distance moved, and 1.9% with duration (Figure 1K-M). There was a high similarity between each cell’s speed and distance correlations (linear regression R^2^ = 0.58, P = 10^-40^) indicating that many of the speed-correlated cells were also distance-correlated. Overall, speed was most strongly correlated with orexin activity (statistics in figure legends). This correlation was evident across behaviour bouts (Figure 1L), irrespective of behaviour class, as well as across averaged metrics within behaviours (Figure 1N).

Head-fixing an animal and allowing it to walk on a treadmill allows both precise measurement of locomotion speed and highly accurate 2-photon recording, so we conducted further studies with a head-fixed apparatus (Figure 2A). 3-dimensional 2-photon imaging of orexin cell activity revealed that during self-initiated movement, while animals were running on a circular treadmill in the dark, orexin neurons fell into five categories based on their activity profiles around the running bout. ‘ON cells’ increased firing, ‘OFF cells’ decreased firing, while ‘down-up cells’ and ‘up-down cells’ had a biphasic firing pattern, and a minority of orexin cells were not significantly modulated during movement (Figure 2A-H; categorization criteria in Methods). Whole-cell recordings combined with epifluorescence imaging in slices confirmed that this range of fluorescence profiles can arise from firing patterns, and that the same orexin cell is biophysically capable of expressing any activity profile (Figure 2I-L). The majority of the recorded orexin cells were up-down (33%) or ON cells (31%), which showed initial increases in spiking activity. Down-up (20%) and OFF (10%) cells, which initially decreased their activity, and nonmodulated cells (6%) made up the remainder (Figure 2M). Each cell type showed a different cross-correlation with treadmill speed (Figure 2N,O). While response onsets were scattered through the time period 2.91 s before to 7.78 s after locomotion start, median response onset time was 0.58 s after movement onset. Many ON and up-down cells had onsets before movement started (Figure 2P,Q). The activity of many ON cells occurred solely during locomotion bouts, whereas many OFF cells were active only outside locomotion (Figure 2R,S; definitions in Methods). These findings suggest that some orexin cells could predict movement before it starts. We assessed this by dividing long recording sessions into two parts, and using the information from the first part to predict when movement would happen in the second part. From the information in the first part, we identified ON cells that had long movement onset lead times and were active solely during movement. A simple threshold-crossing criterion for activity of these cells could be used to detect 80 ± 4 % of imaging frames with movement, and had a false positive rate of only 11 ± 4 % (n = 7 animals). In our highest quality recordings, the weighted average activity vector of these cells predicted movement onset of 75% (9/12) of locomotion bouts 1.55 ± 0.36 s before they occurred (Figure 2T).

**Figure 2.**
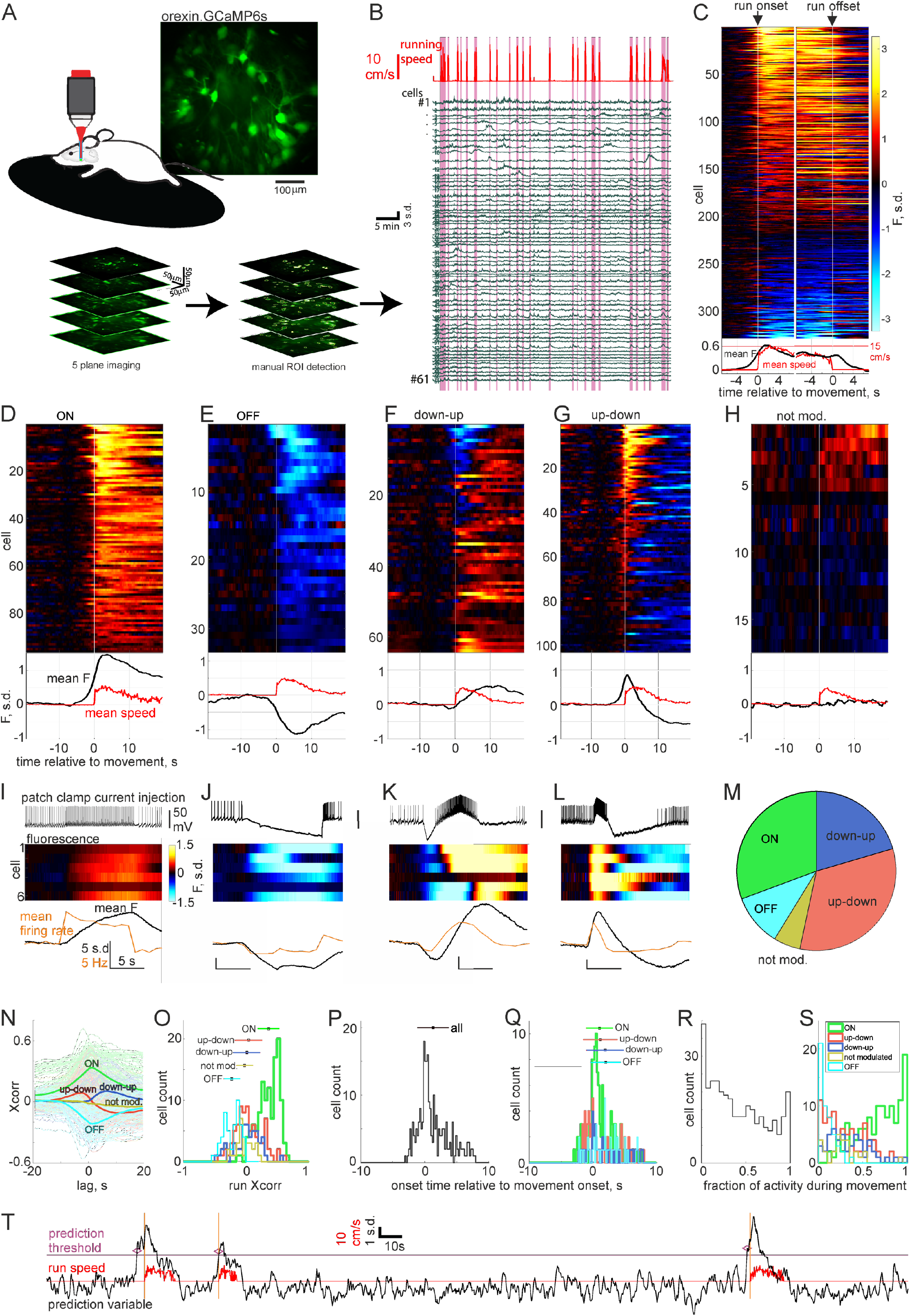
Head-fixed 2-photon imaging of ongoing orexin neuron activity in absence of sensory stimuli. A, Schematic of recording paradigm and example average time projection micrograph (above), and example 5-plane volume imaging with manual ROI detection to average fragments of the same cell across planes (below). B, Example fluorescence activity traces from 61 orexin neurons recorded from one volume and running speed measured from rotation speed of treadmill. C, Averaged activity across self-paced locomotion bouts aligned at locomotion start and end (from 329 cells across 7 animals). Mean fluorescence (black) and run speed (red) across all cells plotted below. D-H, Activity traces from C replotted as subtype groups as defined in methods, aligned to movement start. Scale bar is same as in C. I-L, Data from slice patch clamp recordings combined with Ca-imaging similar to Figure 1C. Cells were recorded in current clamp and injected current waveforms to mimic expected firing profiles in each subtype in D-G (examples shown on top row of each panel). Z-scored fluorescence traces from each cell in middle raster plot and averaged firing rate (orange) and fluorescence (black) across cells in bottom plot. Scale bars are the same for I-L, except for bottom plots where units for scale bars are the same (5 s.d., 5Hz and 5s). M, Proportions of cells in D-H. N, Fluorescence-movement speed cross-correlograms for each cell and averages within subtypes. O, Histogram of fluorescence-movement speed Pearson correlation coefficients. Mean ± s.d. for each subtype plotted at arbitrary heights (ON 0.33 ± 0.17; OFF −0.20 ± 0.13; up-down 0.04 ± 0.20; down-up 0.01 ± 0.20; non mod. −0.02 ± 0.12). P, Histogram of onset time relative to movement start for all cells (mean ± s.d. 1.33 ± 2.40 s). Q, Data from P divided by subtype. Mean ± s.d. for each subtype plotted at arbitrary heights (ON 0.73 ± 1.72 s; OFF 2.03 ± 2.47 s; up-down 1.27 ± 2.77 s; down-up 1.87 ± 2.83 s). R, Fraction of active frames (as defined in methods) that occurred during movement. S, Data from R divided by subtype (ON 0.73 ± 0.23; OFF 0.09 ± 0.08; up-down 0.30 ± 0.23; down-up 0.35 ± 0.22; non mod. 0.28 ± 0.20). T, Prediction variable (black trace) generated from cell fluorescence online plotted against run speed (red). Orange vertical lines denote movement onset, purple horizontal line denotes arbitrary prediction threshold (2 s.d.) and purple diamonds indicate when threshold crossing predicted a run onset.

Since the activity of orexin cells could predict movement, we asked if their activation can elicit locomotion. To probe this, we created a viral construct that directs orexin cells to express the red-shifted channel rhodopsin variant C1V1 (see Methods). This allowed orexin cell firing to be controlled using green laser pulses (Figure 3A-C). Because fidelity of firing decreased at light pulse frequencies above 50 Hz (Figure 3C) and orexin cells are not thought to be naturally active above this frequency in vivo (Lee et al., 2005; Mileykovskiy et al., 2005; Takahashi et al., 2008), we assessed the effects of 2-50 Hz stimulation on locomotion bouts in mice implanted with bilateral light fibers above LH, and transfected with either orexin-C1V1 or orexin-GCaMP6s (as control), while they were head-fixed on a running wheel. Light stimulation elicited running in a frequency dependent manner only in C1V1 expressing mice (Figure 3D,E). The behavioural effect became evident at about 7.5-10 Hz stimulation (Figure 3E), which is within the natural instantaneous firing range of orexin neurons at 0-15 Hz (Lee et al., 2005; Mileykovskiy et al., 2005). In addition to increasing the probability of starting a locomotion bout, increasing stimulation frequency decreased the latency from stimulation to locomotion onset (Figure 3F). Overall, the latencies were distributed around a median of 1.75 s (range 300 – 4880 ms) from stimulus onset which is consistent with the natural latency from orexin ON and up-down cell activation to self-paced locomotion onset (Figure 2Q). These results indicate that orexin neuron activation strongly contributes to rapid initiation of movement.

**Figure 3.**
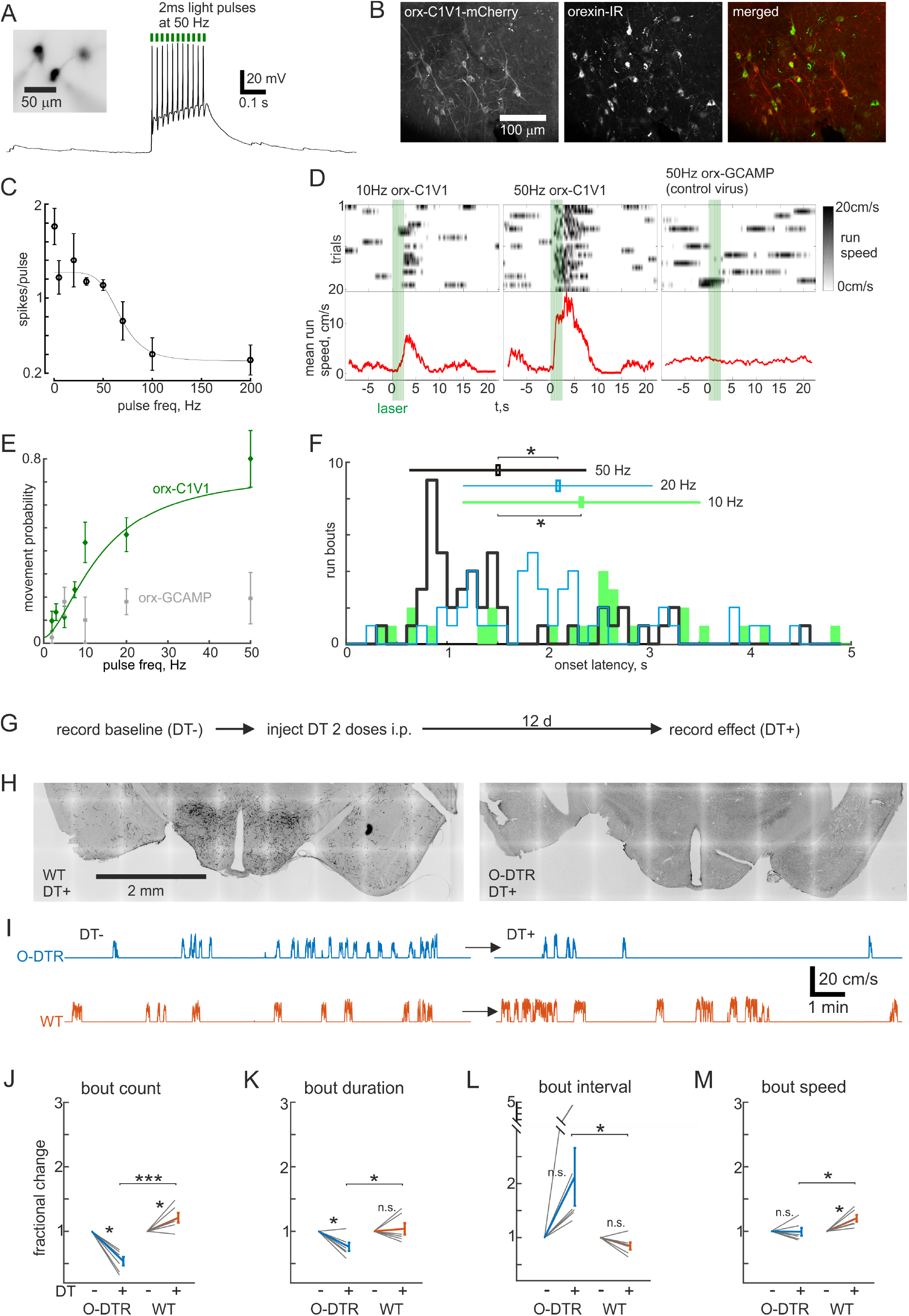
Manipulating orexin neurons controls self-paced movement bidirectionally. A, Micrograph and representative current clamp trace of orexin neuron that expresses C1V1. B, Micrographs showing localization of C1V1-mCherry in orexin immunoreactive (IR) cells. C, Tuning curve of action potential fidelity as a function of light pulse frequency (n=4-6 cells for each data point). Inverse sigmoidal fit explained in methods. D, Treadmill speed across 20 trials and average speed (red) from two head-fixed mice. Mouse with orexin-C1V1 shown with 10 Hz (left) and 50 Hz (middle) light pulses, while mouse with control virus (orx-GCaMP6s) and implanted light fibers shown with 50 Hz light pulses (right). E, Dose response curve of locomotion bout initiation likelihood as a function of light pulse frequency (n=4-6 animals for each data point). F, Histogram of orexin-C1V1 stimulation induced run bout onset latencies from the onset of a 2.5 s light pulse train at the indicated frequencies. Mean ± s.d. for each frequency plotted at arbitrary heights (10Hz 2.33 ± 1.18 s; 20Hz 2.10 ± 0.94 s; 50Hz 1.50 ± 0.88 s, * P<0.01 using the Wilcoxon rank sum test). G, Recording schedule for comparing movement in the same animals with and without orexin cells. H, Representative micrographs of orexin immunoreactivity in wild type (WT) and orexin-diphtheria toxin receptor (O-DTR) brains >12 days after diphtheria toxin injection (DT+). I, Representative treadmill speed recordings from head-fixed O-DTR and WT mice before and after DT administration. J-M, Averaged movement bout statistics normalized to baseline (6 mice/cohort), tested by paired Student’s T-test (*: P<0.05, ***: P<0.0001): J, number of movement bouts during a 16.3 min recording (O-DTR DT+ 54 ± 7 %; WT DT+ 121 ± 8 %); K, duration of bouts (O-DTR DT+ 76 ± 7 %; WT DT+ 104 ± 9 %); L, duration of intervals between bouts (O-DTR DT+ 213 ± 54 %; WT DT+ 84 ± 7 %); M, average speed of bouts (O-DTR DT+ 99 ± 6 %; WT DT+ 120 ± 5 %).

To assess the role of natural orexin cell activity in self-initiated locomotion, we next performed three loss-of-function experiments. First, we ablated orexin neurons using a mouse line expressing the human diphtheria toxin receptor in orexin neurons (O-DTR) (González et al., 2016b). When these mice were injected with a low dose of diphtheria toxin (DT), their orexin neurons were deleted (3.6 ± 1.8 % remaining, n = 6 animals) within 12 days, while DT-injected wild-type control mice maintained their orexin neurons (Figure 3G,H). We compared statistics of self-initiated locomotion in the dark before and after DT administration to O-DTR and wild-type mice (Figure 3I-M). After orexin neuron deletion, mice ran significantly fewer bouts than prior to deletion, or than in wild-type controls (Figure 3J). The bouts were also significantly shorter, by 24 ± 7 *%* compared to before deletion (Figure 3K). Additionally, in orexin-neuron-ablated mice stationary intervals between runs were longer and locomotion speed was lower than in wild-type controls (Figure 3L,M). These results suggest that orexin neurons regulate self-generated movement initiation, maintenance and speed.

Secondly, to complement this chronic manipulation of orexin neurons, and to selectively erase activity increases (Figure 2), we created a viral construct that directs orexin neurons to express the red-shifted inhibitory opsin ArchT (see Methods). This allowed us to hyperpolarize orexin neurons with green laser light by 26.7 ± 6.7 mV (n = 8 cells, Figure 4A). With this temporally controllable, reversible loss-of-function approach we could interleave laser-on and laser-off trials and compare to control mice injected with the GCaMP6s virus. To make comparisons across trials consistent, we waited for mice to be stationary for 10 s before either turning the laser on for 30 s to inhibit orexin neurons, or recording a catch trial (laser off). We quantified the movement bouts during this 30 s period. Orexin neuron inhibition decreased the number of self-initiated locomotion bouts (Figure 4C-E), but did not affect bout duration, latency to run onset during the assay period, or speed of the run bouts (Figure 4F-H). In the third loss-of-function experiment we asked what would happen if orexin cells were inhibited after movement onset. We turned the laser on for 10 s as soon as a run onset was detected (~0.1-0.5 s delay) with interleaved catch trials (laser off) with at least 30 s intervals between consecutive trials. There was no effect on duration or speed of runs, suggesting that orexin cell activity after locomotion initiation is not necessary for maintenance of locomotion. Taken together, these acute optogenetic loss-of-function experiments indicate that the natural activation of orexin cells initiates self-generated locomotion, but does not modulate the parameters of locomotion bouts once they have started. Unlike optogenetic inhibition, orexin neuron deletion also decreased run duration and speed (compare Figures 3K,M and 4F,H), which suggests that the inhibitory components of OFF, up-down and down-up cells might control these parameters (see Discussion).

**Figure 4.**
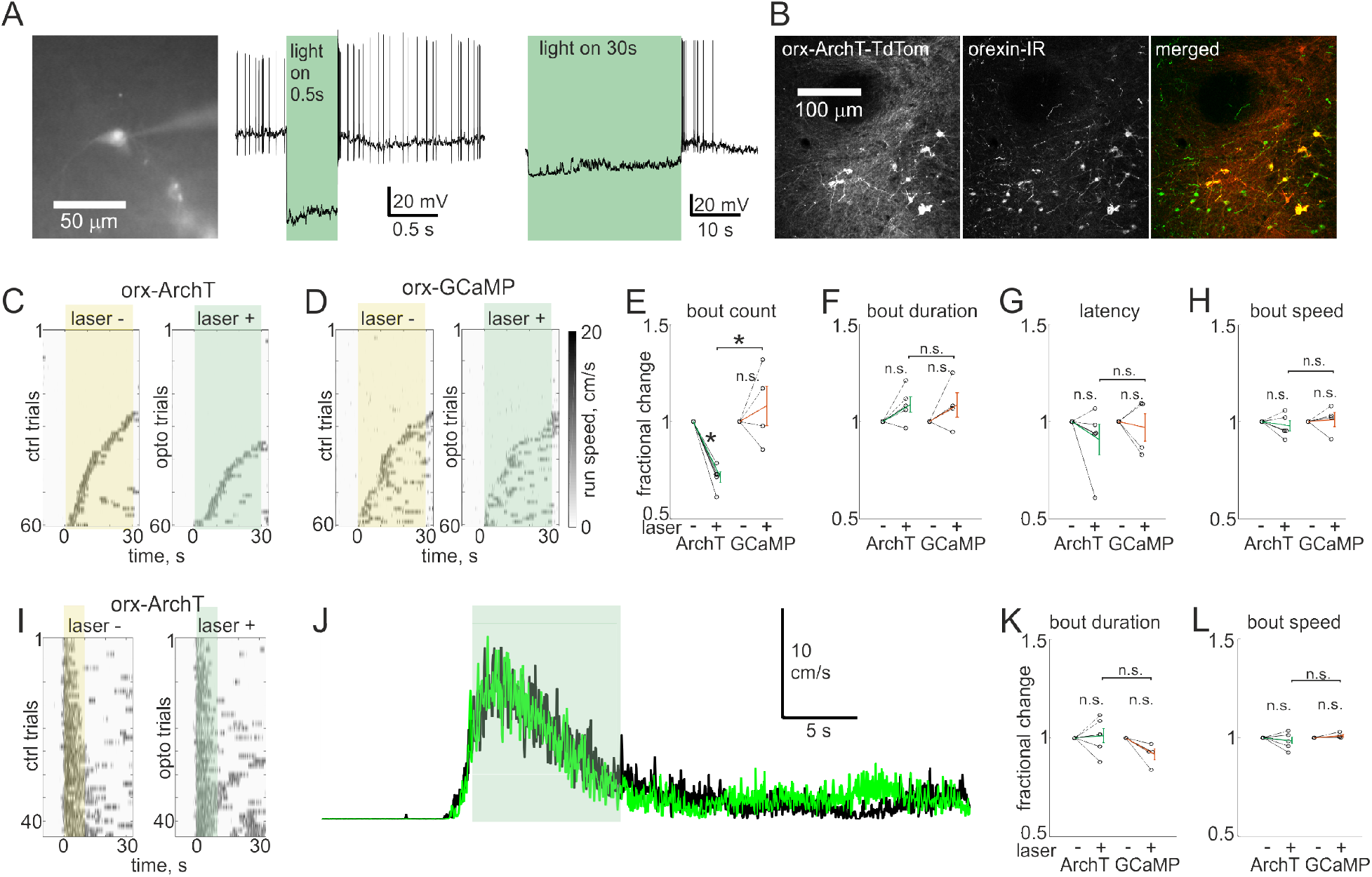
Acute inhibition of orexin neuron activation decreases only movement initiation. A, Representative micrograph and current clamp recordings of ArchT expressing orexin neuron. Representative micrographs of ArchT-TdTomato expression in orexin immunoreactive (IR) neurons. C-D, Example recordings of treadmill speed during self-initiated movement in orexin-ArchT and control (orexin-GCaMP6s with light fiber implants) animals. E-H, Averaged movement bout statistics normalized to baseline from each mouse (5 ArchT mice and 4 control mice), tested by paired Student’s T-test (*: P<0.01). E, number of movement bouts (ArchT 71 ± 3 %; GCaMP 108 ± 10 %); F, duration of bouts (ArchT 109 ± 4 %; GCaMP 109 ± 6 %); G, latency from laser pulse to run bout (ArchT 91 ± 8 %; GCaMP 97 ± 7 %); H, average speed of bouts (ArchT 98 ± 3 %; GCaMP 101 ± 4 %). I, Example recordings of treadmill speed during self-paced running where laser was turned on after movement was detected (on laser+ trials). Scale same as in D. J, Average running speed across trials in I, green is from laser+ and black from laser-(catch) trials. K-L, Averaged movement bout statistics normalized to baseline from each mouse (5 ArchT mice and 4 control mice), tested by paired Student’s T-test (n.s.: P>0.05). K, duration of bouts (ArchT 101 ± 4 %; GCaMP 92 ± 3 %); L, average speed of bouts (ArchT 97 ± 1 %; GCaMP 101 ± 1 %).

Given that the main effect of the loss-of-function manipulations was decreased initiation of self-generated movement, we next wanted to inspect the role of orexin neurons in sensory-evoked movement. During freely-moving recordings, we noticed many events that looked like sensory responses, because they coincided with tactile, visual or olfactory stimuli (Figure 5A). However, it was unclear whether these responses were in fact due to something else, such as movement. Therefore, we presented head-fixed the mice with sensory stimuli and only analysed responses when the subject did not move during the trial (Figure 5B). We also recorded the activity profiles of the same cells during self-initiated locomotion and compared responses across modalities and cell types (Figure 5C-F). All cell types responded with transiently increased activity to visual (blue LED flash), tactile (airpuff to left side abdomen) and olfactory (amylacetate or female urine) stimuli. In all recordings, we estimated the distinction between noise and true response by interleaved presentation of a null stimulus (see Methods). These null responses allowed us to delineate thresholds of detection throughout our analysis (null responses fell within dashed black boxes in Figure 5C-E,G-I). This revealed only two cell-stimulus pairs with inhibitory responses (points on negative side of black boxes in Figures 5C-E). In total 62.0% of orexin cells had excitatory responses to at least one modality, and of these 58.5% responded to more than one modality and 17.0% to all three modalities (Figure 5F). Since 57.8% of responsive cells had visual, 33.3% tactile and 84.4% had olfactory responses, the proportions of bimodal and trimodal cells were consistent with chance overlap. This suggests that orexin cells are not tuned to a particular sensory modality. We also captured some self-initiated locomotion bouts during epochs with no sensory stimulus (Figure 5B right panel). These data showed that magnitude of peak locomotion modulation did not correlate with the size of sensory responses but the magnitude of sensory responses in ON cells was significantly greater than in OFF cells (P<0.001 using the Wilcoxon rank sum test, Figure 5G-I), suggesting that sensory evoked movements could potentially be elicited through activation of ON cells. Indeed, the ON cell population had the highest proportion of responsive cells (71% responded to at least one stimulus modality), while OFF cells had the lowest (33%), and up-down 67%, down-up 50% and non-modulated 33%.

**Figure 5.**
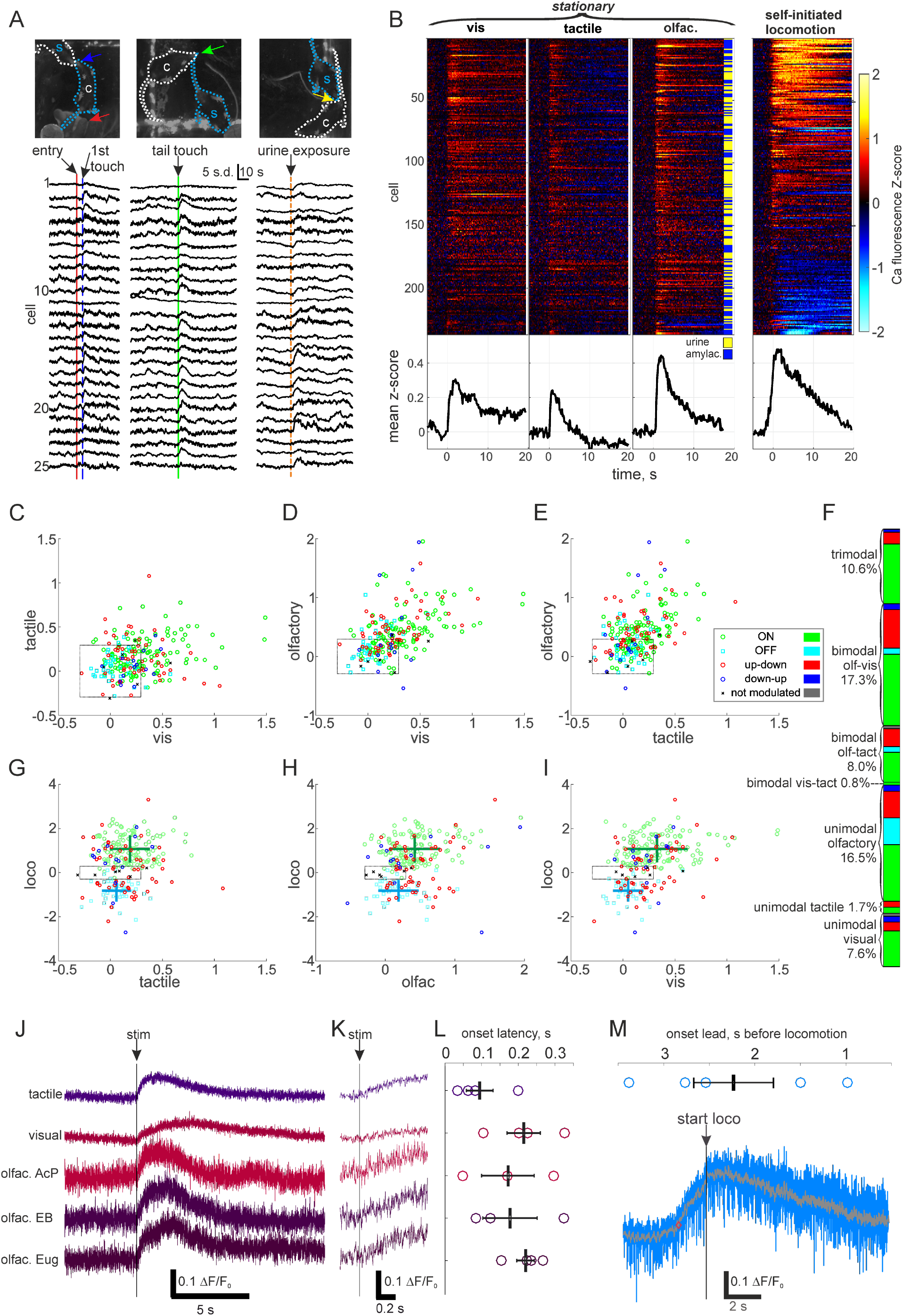
Sensory responses in orexin cells. A, Example natural sensory responses to social stimuli in orexin cells recorded with miniature endoscope in freely moving male subject (s, blue dashed outline) while it was in the arena with a female conspecific (c, white dashed outline). Arrows indicate locations of natural stimuli. In left panel conspecific enters arena from handler’s hand (red arrow) and moves to touch snouts with subject (blue arrow). In middle panel conspecific investigates subjects tail, touching it (green arrow) and causing subject to stop feeding and turn around. In right panel subject is investigating conspecific’s rear when conspecific begins to urinate (yellow arrow). B, Head-fixed sensory responses from 237 cells from 4 mice, averaged from 5-11 trials where mice remained stationary (three panels from left). Vis=visual, blue LED flash; tactile, 30 psi pressure pulse applied to left side abdomen; olfac.=olfactory, odour pulse (rapid valve switch between empty vial and odour vial with no change in pressure) to snout containing amylacetate (amylac.) or urine pooled from females in different cages. Right panel contains averaged locomotion activity profiles aligned to self-initiated movement onset from each cell from 5-22 locomotion bouts outside sensory stimulation trials. Black traces at the bottom of each cell raster are averages across all cells. C-E, Scatter plots of response amplitudes (mean signal from 1-4 s after stimulus) of all cells to two modalities in each plot, coloured by the locomotion subtype (legend in E). Dashed black boxes denote the extrema of null-responses (quantified response amplitude to a 77dB 10kHz sound stimulus that did not cause a detectable response), i.e., only responses outside the box should be considered meaningful. F, Proportions of responsive cells out of all cells coloured by locomotion subtype and grouped by the number of response modalities. G-I, Scatter plots of response amplitudes and locomotion profile averages (see methods). Legend as in E. Green and cyan crosses denote mean ± s.d. for ON and OFF cell populations respectively (sensory responses: G, ON 0.19 ± 0.19, OFF 0.06 ± 0.14; H, ON 0.42 ± 0.36, OFF 0.19 ± 0.29; I, ON 0.33 ± 0.29, OFF 0.05 ± 0.15) which were significantly different for all modalities (P<0.001 Wilcoxon rank sum test). J, Fiber photometry responses of the orexin population to tactile, visual and three different olfactory stimuli (AcP, acetophenol; EB, ethylbutyrate; Eug, eugenol). K, Expanded view from J. L, Quantified latency from stimulus onset to response onset tactile (95 ± 36 ms), visual (172 ± 72 ms) and three different olfactory stimuli (acetophenol, 178 ± 74 ms; ethylbutyrate, 220 ± 24 ms; eugenol, 214 ± 45 ms) in averaged traces (5-20 trials) for 4 mice for all stimuli except AcP and EB from 3 mice. M, Onset lead delay from fiber photometry signal increase to movement onset (2230 ± 440 ms) from 11-28 locomotion bouts from 5 mice. Blue background trace is unfiltered and black trace is smoothed with a 100-sample moving average.

We used fiber photometry of average signal across GCaMP6s expressing orexin cells to quantify sensory latencies, because this allowed increasing sampling rate to 500 Hz. Sensory responses started on average 176 ± 22 ms after stimulus (range 34 – 324 ms), while orexin population activity increased above baseline 2230 ± 440 ms before self-initiated locomotion onset (Figure 5J-K). This timing is consistent with the possibility that sensory activation of orexin cells could lead to locomotion.

We probed orexin neuron-mediated sensory-evoked movement further by analysing orexin neuron activity during stimulation trials where the stimulus evoked movement (Figure 6A-C). A gentle airpuff to the base of the tail induced locomotion in 33% (16/48) of trials, and female urine odour induced locomotion in 25% (12/48) of trials across four animals. When sensory stimulation evoked locomotion, the overall orexin cell responses were bigger than when mice remained stationary during stimulus presentation (Figure 6C,D). To compare orexin cell activity during sensory-evoked movement to their activity profiles during self-initiated locomotion, we aligned the sensory stimulus trials where movement was triggered to locomotion onset (Figure 6B). This showed that time-courses were comparable between the conditions, while sensory-evoked movement was coupled with higher amplitudes than the activity profiles during self-initiated locomotion. ON cells in particular were significantly more active during sensory-evoked movement than during sensory responses while stationary (Figure 6D). Therefore, in order to examine objective predictive information about future movement in this orexin subpopulation, we used a receiver operating characteristic (ROC) analysis to objectively assess the evolution of the ROC AUC (area under the curve) predictor in ON cells from sensory stimulation until locomotion onset (Figure 6E). This showed that an ideal observer could use ON cell activity to predict the decision to move 600 ms after the sensory stimulus with 73% accuracy. At the time point where half of the evoked movement bouts had begun (2 s after stimulus onset), ON cell activity predicted the future movement onsets with 93% accuracy (instantaneous ROC AUC values in Figure 6E were only calculated from trials where movement had not yet begun). These analyses indicate that ON cells are highly activated before sensory-evoked movement, and can predict it.

**Figure 6.**
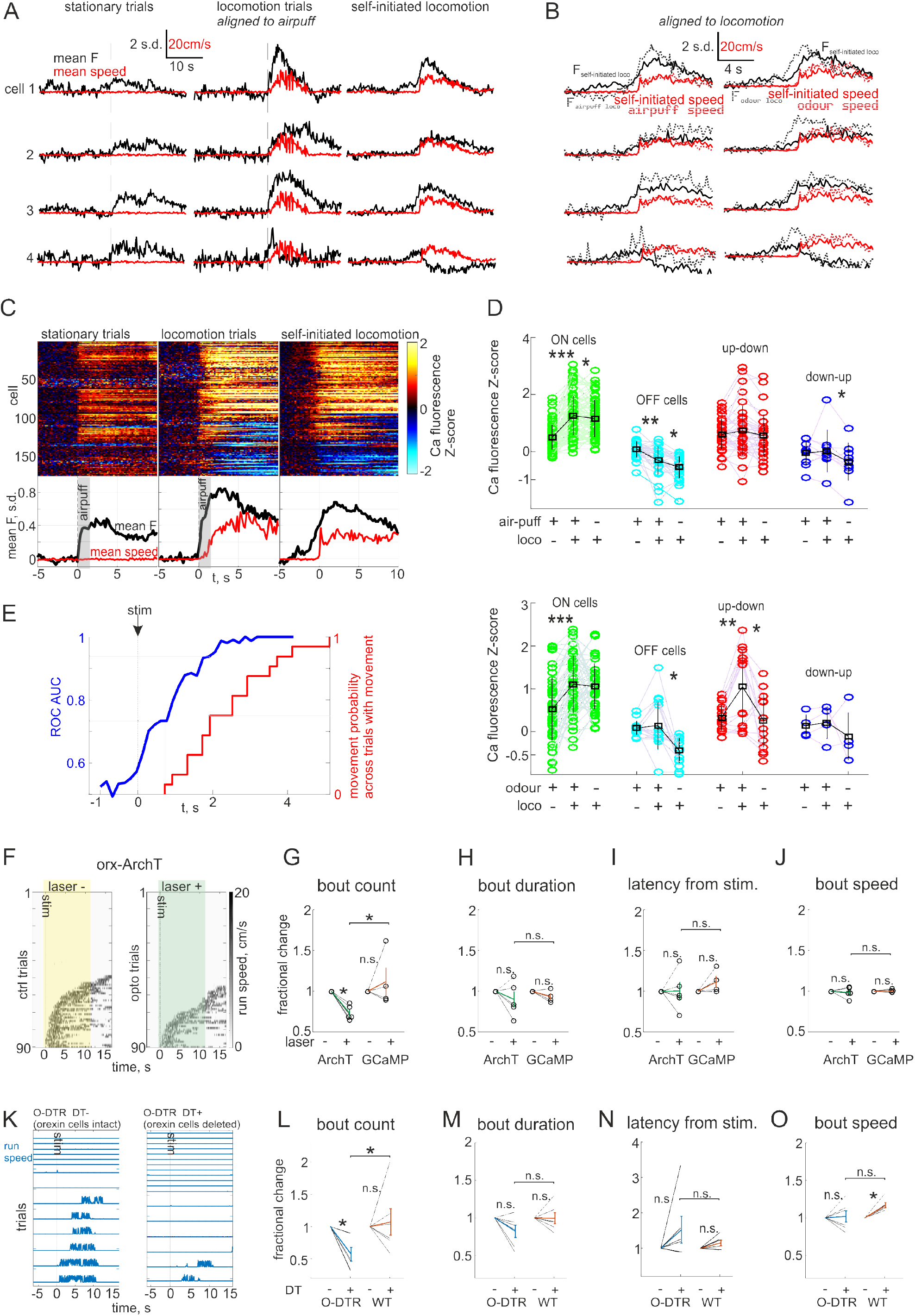
Orexin cells mediate a rapid sensorimotor transformation. A, Example cells responding to tactile stimulus (2 s airpuff 15 psi to base of tail) while mouse was stationary (left) and when the airpuff triggered movement (middle). Also shown activity profiles during self-paced movement without sensory stimuli (right). B, Expanded view of self-paced traces from A (black and red) and stimulus triggered movement (magenta, airpuff on left; social odour on right, consisting of female urine odour presentation for 0.5 s) and response (blue, airpuff on left; social odour on right) aligned to the onset of movement. C, Activity of all recorded cells in response to airpuff while mice were stationary (left, averaged across) and when movement was triggered (middle). Also shown activity profiles during self-paced movement without sensory stimuli (right). Bottom black traces are averages across all cells and red traces movement speed. D, Mean responses measured from an analysis window 1-4 s after sensory stimulus onset (or locomotion onset for self-paced locomotion) in airpuff (above, mean ± s.d. values in black: ON 0.50 ± 0.42, 1.27 ± 0.67, 1.16 ± 0.63; OFF 0.09 ± 0.29, −0.31 ± 0.50, −0.54 ± 0.40; up-down 0.61 ± 0.53, 0.73 ± 0.87, 0.56 ± 0.73; down-up −0.05 ± 0.43, 0.02 ± 0.73, −0.36 ± 0.65) and odour presentation (below, mean ± s.d. values in black: ON 0.53 ± 0.73, 1.11 ± 0.67, 1.06 ± 0.54; OFF 0.08 ± 0.16, 0.14 ± 0.56, −0.44 ± 0.30; up-down 0.33 ± 0.30, 1.05 ± 0.87, 0.26 ± 0.62; down-up 0.14 ± 0.25, 0.21 ± 0.37, −0.11 ± 0.55). * P<0.025, ** P<0.001, *** P<10^-8^. E, Evolution of area under the curve receiver operating characteristic calculated from ON cell activity data from airpuff stimulation trials where locomotion had not yet started (blue). Red plot is the cumulative probability distribution of locomotion onsets from trials with movement (16/48). Grid-lines indicate stimulus onset, first locomotion bout onset and half of locomotion bouts started. F, Example recordings of treadmill speed during sensory evoked running in head-fixed orexin-ArchT animals with laser turned on 0.5 s before stimulus (left) and without laser (right). G-J, Averaged movement bout statistics normalized to baseline from each mouse (5 ArchT mice and 4 control mice), tested by paired Student’s T-test (*: P<0.05). G, number of movement bouts (ArchT 73 ± 4 %; GCaMP 112 ± 17 %); H, duration of bouts (ArchT 90 ± 10 %; GCaMP 93 ± 4 %); I, latency from sensory stimulus to run bout (ArchT 100 ± 11 %; GCaMP 111 ± 7 %); H, average speed of bouts (ArchT 99 ± 3 %; GCaMP 101 ± 1 %). K, Representative treadmill speed recordings during sensory evoked running from head-fixed O-DTR mice after (left) and before (right) DT administration. L-O, Averaged movement bout statistics normalized to baseline from each cohort (6 mice/cohort), tested by paired Student’s T-test (*: P<0.05). L, number of movement bouts (O-DTR DT+ 50 ± 10 %; WT DT+ 107 ± 21 %); M, duration of bouts (O-DTR DT+ 88 ± 8 %; WT DT+ 100 ± 7 %); N, latency from sensory stimulus to run bout (O-DTR DT+ 165 ± 44 %; WT DT+ 115 ± 8 %); O, average speed of bouts (O-DTR DT+ 100 ± 9 %; WT DT+ 117 ± 3 %).

Finally, in order to examine if there is a causal link between orexin cell activation and sensory-evoked movement, we tested whether erasing orexin cell activation by ArchT mediated hyperpolarization would block this sensorimotor transformation. When LH orexin-ArchT cells were bilaterally inhibited from 0.05 s before to 12 s after a sensory stimulus, significantly fewer locomotion bouts were elicited than in interleaved control stimulation trials, and this difference was not seen in control virus (orexin-GCaMP) injected mice (Figure 6F,G). However, this peri-sensory orexin cell opto-silencing elicited no change in duration, latency or speed of the movement (Figure 6H-J). This indicates that excitatory components of orexin cell responses, like ON cell activation, are critical for normal sensorimotor transformations. To determine if the initial inhibitory responses in OFF and down-up cells might play a role in sensorimotor transformations, we further inspected the role of all orexin cells in sensory-evoked movement by deleting them in O-DTR mice. The effect of O-DTR DT+ orexin neuron deletion was a decrease in occurrence of sensory-evoked movement (Figure 6K-O), similar to the ArchT experiment. This suggests that the activation rather than inactivation of orexin neuron subpopulations accounts for their role in sensory-evoked movement. Overall, these results demonstrate that orexin neurons selectively affect sensory-evoked movement initiation rather than maintenance, reaction time or speed.

## Discussion

Our data provide strong evidence for a key role of orexin neuron activity in millisecond-scale sensorimotor transformations (Figures 5 and 6) as well as self-initiated movement (Figures 2–4) and diverse mobile behaviours (Figure 1). Previous work has demonstrated that orexin neurons have sensory responses (González et al., 2016a; Hassani et al., 2016; Inutsuka et al., 2016; Mileykovskiy et al., 2005), and here we have demonstrated that these responses have a function in coupling sensation to action. To our knowledge, this is the first hypothalamic neuronal population shown to mediate a rapid sensorimotor transformation, similar to those seen in the neocortex (Ferezou et al., 2007; Svoboda and Li, 2018). The broad stimulus receptivity (Figure 5) and robust activation during mobile behaviours (Figure 1) indicate that orexin neurons have a broader and more immediate influence on awake behaviour than previous literature suggests (Mahler et al., 2014; Tyree et al., 2018).

Based on their activity profiles during self-initiated movement, orexin neurons can be functionally divided into several subclasses. Of the movement-related subtypes, ON cells carried clear predictive information about future movements and were robustly activated when sensory stimuli elicited movement (Figure 6D,E). By comparing two loss-of-function manipulations – acute reversible hyperpolarization and chronic deletion of orexin neurons – we were able to distinguish between the effects of disrupting the excitatory components (with ArchT), or both excitatory and inhibitory components (with O-DTR) of orexin cell responses. This indicated that the early excitatory components, i.e. activity of ON and up-down orexin cells, are critical for initiating self-generated and sensory-evoked movement, while the inhibitory components may influence the duration and speed of internally generated movement. Alternatively, the effects of the chronic orexin-DTR cell deletion could be due to compensatory mechanisms, or due to higher magnitude and penetrance of the manipulation (see Methods).

Orexin neurons were previously thought to homeostatically control overall locomotor activity in response to slow manipulations such as fasting (Hu et al., 2015; Yamanaka et al., 2003). Our results demonstrate that orexin neurons are also critically important for sub-second rapid regulation of locomotion and sensorimotor processing. The shortest timescale we captured here was 34 – 324 ms from sensory stimulus to orexin cell activity increase (Figure 5K). This activity predicted sensory-evoked movement some 600 – 4600 ms after stimulus onset (Figure 6E).

Exogenous activation of orexin cells with C1V1 led to movement in 300 – 4880 ms (Figure 3F), while internally generated activation of orexin cells could precede movement by up to 3380 ms (Figure 5L). It is therefore clear that hypothalamic neurons, despite their paradigmatic position as slow modulators of whole body physiology, have a rapid effect on motor performance.

The sensory response latencies of orexin cells (34 – 324 ms, average 176 ms) were on par with our measurement of delays recorded from action potential burst to GCaMP6s fluorescence increase (31 – 288 ms, average 149 ms; Figure 1C). The difference of these averages (23 ms) implies that the sensory responses are mediated via a small number of synaptic contacts similar to classical cortical sensory responses (Allen et al., 2017; Ferezou et al., 2007; Mohajerani et al., 2013). The sensory signals could thus arise through various multi-synaptic pathways, for example, from cortex, central gray or amygdala. In principle, it is possible that the signals in freely-moving recordings (Figure 1) are communicated from cortical neurons tuned to spatial location, head direction or vestibular input (Moser et al., 2017; Vélez-Fort et al., 2018). However, this is made less likely by the fact that the signals were maintained while the head was fixed in space (Figures 2 and 6).

Our work places orexin neurons into a growing diagram of movement controlling neurons across the brain (Arber and Costa, 2018; Svoboda and Li, 2018). In particular, orexin neurons initiate locomotion, consistent with ‘stepping’ induced by electrical stimulation of LH in anaesthetized animals (Jordan, 1998; Sinnamon, 1993). A recent study demonstrated similar properties in substantia nigra pars compacta dopamine neurons during self-initiated locomotion; this population had heterogeneous activity profiles during movement with some cells turning off and most being activated, and optogenetic manipulation revealed a highly similar phenotype to what we have shown here for orexin neurons (da Silva et al., 2018). Similarly, classical movement-control neurons in the basal ganglia activate predictably before self-initiated movement (Cui et al., 2013). Thus, we find that orexin cells closely resemble classical subcortical motor control neurons in the midbrain and striatum. As orexin neurons are in a unique position to affect sympathetic outflow (Bonnavion et al., 2015; Kerman et al., 2007) their motor activation could potentially function to prepare the body for movement, for example, by increasing heart-rate and respiration.

Future work should inspect synaptic interactions between orexin cells and other movement controlling neurons to elucidate how these populations orchestrate decisions to move. It should also assess whether orexin cell movement-subtypes correlate with known electrophysiological, molecular or anatomical subtypes (Apergis-Schoute et al., 2015; Iyer et al., 2018; Mickelsen et al., 2019, 2017; Schöne et al., 2011). It has been hypothesized that orexin neurons promote exploratory activity, hunger-driven food-seeking or any motivated behaviour (Mahler et al., 2014; Tyree et al., 2018; Yamanaka et al., 2003). Future work should therefore also inspect whether orexin activation is specifically associated with movement toward a particular goal, given the option for an animal to move in order to achieve different outcomes. However, pending a more detailed assessment, the present results suggest that orexin neurons would control locomotor movement regardless of the goal, as expected from circuits underlying basal motivation or motor control required for all movement.

## Methods

### Animals

Animal handling and experimentation was approved by the UK government (Home Office) and by Institutional Animal Welfare Ethical Review Panel. Animals of both sexes, aged 30-60 days at the beginning of the procedures were used and were housed in a controlled environment on a reversed 12h light-dark cycle with food and water ad libitum. WT C57BL6 mice were obtained originally from the Jackson Laboratory. Mice expressing the human diphtheria toxin receptor in orexin cells (O-DTR mice) were generated as described before (González et al., 2016b) and cross bred with WT mice. The specific deletion of orexin neurons with DT in this strain has been documented before (González et al., 2016b) and was confirmed in the experimental animals in this study (see Figure 3H). O-DTR mice had 3.6 ± 1.8 % orexin neurons remaining compared to WT mice after DT (cell counts from 6 mice, 3 bilateral sections from each mouse: 2268 orexin neurons in WT DT+ and 82 in O-DTR DT+). For fiber photometry experiments in Figure 5 J-L, orexin-cre mice (Schöne et al., 2012) were used.

### Surgery: Virus injections

Two to three weeks prior to optical device implantation, WT mice were injected stereotactically with AAV1-hORX.GCaMP6s (prepared by Penn vector core, PA, USA), AAV1-hORX.C1V1(t/s).mCherry or AAV1-hORX.ArchT.TdTomato (prepared by Vigene Biosciences, MD, USA). The GCaMP6s virus was generated as described before (González et al., 2016b), and the other two were cloned using the same plasmid and plasmids acquired from Addgene. Using the orx.GCaMP6s virus, GCaMP6s was expressed with 97.4 ± 1.0 % specificity and 65.8 ± 3.7 % penetrance in orexin neurons (381 ± 73 orexin+, 242 ± 40 GCaMP6s/orexin+ and 6 ± 2 GCaMP6s/orexin-cells counted from 4 animals, Figure 1G). The mean delay from onset of a spiking train to detected fluorescence increase was 149 ± 30 ms (n=6 cells, range 31 – 288 ms, Figure 1D). For experiments in Figure 5J-L, orexin-cre mice were injected in the left hemispheres with AAV9-CAG.Flex.GCaMP6s.WPRE.SV40 (prepared by Penn vector core, PA, USA) and expression has been characterized before (González et al., 2016b, 2016a).

Using the orx.C1V1.mCherry virus, C1V1.mCherry was expressed with 96.0 ± 0.2 % specificity and 54.8 ± 3.4 % penetrance in orexin neurons (224 ± 36 orexin+, 119 ± 13 mCherry/orexin+ and 6 ± 3 mCherry/orexin-cells counted from 4 animals, Figure 3B). Using the orx.ArchT.TdTomato virus, ArchT.TdTomato was expressed with 99.1 ± 0.4 % specificity and 59.1 ± 3.3 % penetrance in orexin neurons (289 ± 92 orexin+, 161 ± 38 TdTomato/orexin+ and 1 ± 1 TdTomato/orexin-cells counted from 5 animals, Figure 4B). For surgery, mice were anesthetized with isoflurane, the scalp was infiltrated with lidocaine, opened, and a 0.2 mm craniotomy was drilled at 0.9 mm lateral, 1.4 mm posterior from Bregma (in the right hemisphere for imaging experiments and bilaterally for optogenetic manipulation and control experiments). A pulled glass injection needle was used to inject 100-400 nl of virus 5.4 mm deep in the brain at a rate of 50 nl/min. After removal of the injection needle, the scalp was sutured and animals received 5 mg/kg carprofen injections for two days as post-operative pain medication.

### Surgery: optical device implantation

Two weeks after viral delivery mice were anaesthetized with isoflurane, the scalp infiltrated with lidocaine, and opened. A custom-made aluminium head plate was attached to the skull using three skull screws and dental cement (Metabond). A 0.8 mm diameter hole was drilled at the same position(s) as in virus injection surgery and the optical device was slowly (150μm/min) lowered to a depth of 5.1 mm using an automated Luigs & Neumann micromanipulator controlled from Matlab. Lens placements in Supplementary figure 1. For imaging experiments, the optical device was a 0.39 NA, 7.3 mm long, 0.6 mm diameter cylindrical graded refractive index (GRIN) lens (Grintech). For fiber photometry experiments in Figure 5J-L, the optical device was a 200 μm core fiber optic cannula (Doric MFC_200/260-0.37_50mm_MF2.5(7.5mm)_FLT, or ThorLabs CFML12U-20, NA 0.39). For optogenetic experiments (and GCaMP controls in optogenetic experiments), the optical device consisted of two 0.39 NA, 10 mm long, 0.2 mm diameter optic fiber cannulae. After lowering optical devices into the LH, they were cemented onto the skull and the implant was coated with black dental cement and painted with 3-4 layers of black nail polish to keep scattered light from entering the environment.

The mice received a single dose of 0.6mg/kg dexamethasone and 5mg/kg carprofen injections for two days as post-operative pain and anti-inflammatory medication. After at least two weeks of recovery, mice were trained for head-fixed awake experiments in 5-10 sessions of increasing length on the experimental running-wheel apparatus or for freely moving miniature endoscope recordings in a similar incremental regimen of wearing the head-mounted miniature microscope (~2g weight) in the test arena (Figure 1F).

### Freely moving Ca^+2^ imaging

Freely moving Ca imaging was performed with a head-mounted miniature microscope (Inscopix) which was attached to the animal’s head without anaesthesia to a magnetic baseplate that had previously been cemented at the correct focal height on top of the implant. The subject was then placed into the plexiglass-floored arena which it had already spent time in and recording was begun after 10 minutes. A typical recording session lasted 30 minutes during which time a familiar opposite-sex conspecific was introduced by a gloved handler (conspecific was supported on palm rather than held by tail during entry), and sometimes (rarely) the handler’s gloved hand would ‘chase’ the subject. Most of each recording was simply observation of the subject and conspecific in a familiar arena with wet food available ad libitum. The arena was lit with red LED lighting and was covered from view in a dark, quiet room and a silent fume extractor pipe was located 50 cm above the arena to eliminate background odours. Ca^+2^ imaging was performed at a frame rate of 10 frames/s. Behaviour tracking was done with a CCD camera (Lumenera Infinity) below the arena acquiring at 60 frames/s and synchronized with Calcium imaging using an LED mounted next to the arena and facing the behaviour camera. The LED was driven to flash on every acquired Calcium imaging frame with a TTL pulse from the microscope DAC board. Behaviour analysis was done in ImageJ by manually classifying behaviours and measuring distance and time travelled during each behaviour bout.

### Head-fixed two-photon Ca^+2^ imaging

Mice were imaged for 0.5-1 h head-fixed on a running disk allowing the animal to move or remain stationary ad libitum. Imaging was performed either in a dark room with red background lighting (Figures 2 and 5) or with a 21-inch flat screen monitor located 15 cm away from the animal’s eyes and displaying a low luminosity picture of a natural forest scene (Figure 6). Throughout imaging sessions, we recorded the movement of the running disk via a custom-built infra-red optical sensor which was sensitive to small movements and encoded speed linearly as the differential of the optical signal. Locomotion epochs were defined as the time when the mouse moved >1 cm/s. A breathing sensor was placed near the right nostril to monitor normal breathing.

Changes in GCaMP6s fluorescence were imaged with a custom electro-tunable lens equipped resonant/galvanometer scan head two-photon microscope (INSS) and a femtosecond-pulsed mode-locked Ti:sapphire laser (Spectra-physics Mai Tai Deepsee 2) at 950 nm through a 20 x (0.45 NA, Olympus) air-IR objective at 30.1 frames/s with 512 x 512 pixel resolution using custom Labview software. With 5-plane imaging using the electro-tunable lens volume rate was 5.1 volumes/s. Images were obtained with a 510/80 nm band-pass emission filter.

Visual stimuli were delivered with a blue LED positioned at eye level 10 cm away and visible to both eyes. Tactile stimuli were delivered with a fast TTL driven valve device (picospritzer) through a through a fan shaped nozzle (15 mm length, 3 mm width, which formed a air stream approximating a fan with 60° angle) so that the geometry of the airpuff impinging on the animal was insensitive to small postural changes. Tactile stimuli were either 0.2 s pulses of 30 psi pressure delivered to the left side lower abdomen from the side as a vertical fan pattern (Figure 5) or 2 s pulses of 15 psi pressure delivered to the base of the tail from the back as a horizontal fan pattern (Figure 6). Olfactory stimuli were delivered with a custom TTL driven solenoid valve switching device, covered in a sound-proof case, which delivered a constant 1 psi stream of air through an empty vial to the animal’s snout from 1 cm distance. The device created odour pulses by switching valves rapidly to direct the air through an odour vial containing amylacetate, female urine (~0.5-1ml collected fresh from 5 separately housed female mice), ethylbutyrate, acetophenol or eugenol. Odour pulses and background odours were cleared by a constant-suction high-capacity fume extrusion pipe (8 cm diameter) located 5 cm in front of the animal which was there during all recordings. Odour delivery and lack of pressure changes was confirmed by the experimenter. The null stimulus in Figure 5 was an auditory stimulus delivered through speakers located 15 cm in front of the animal and driven by a 10kHz pure tone clip played from Matlab using the psychophysics tool box. The auditory stimulus intensity was 77 dB and background noise on the microscope was 65 dB, and it did not generate sensory responses in orexin neurons.

The presentation of sensory stimuli was synchronized with image acquisition using custom routines in Matlab to generate timing triggers. Within Matlab, a National Instruments USB-6008 DAQ board was used to output TTL triggers and count imaging frames. Stimuli were presented in blocks consisting of a visual, tactile, auditory and 1-3 odours presented for 0.2 s (Figure 5) or tactile and an female urine odour presented for 2s and 0.5 s respectively (Figure 6). Stimulus interval was 15 s plus a randomly selected 0.1-15 s re-randomized before every stimulus presentation (Figure 5) or 30 s plus a randomly selected 0.1-30 s (Figure 6). In addition, order of the stimuli was randomized for each block. Before every experiment mice were allowed to habituate to head-fixation and stimulus presentation for 5-10 min.

### Head-fixed Ca^+2^ fiber photometry

GCaMP6s was excited with a blue LED (Prizmatix UHP-LED-460, 460 nm, ThorLabs M470F3, 470 nm, Figure 5M) or a 473 nm laser (Becker & Hickl, Figure 5J-L) and emission was sampled at 500 Hz with a photoreceiver (Newport 2151 or Becker & Hickl HPM-CON-2) through a fiber-connectorized GFP filter cube (Doric, FMC_GFP_FC). Optic fibers were from ThorLabs (multimodal, FT200EMT, 200 μm core diameter, 0.39 NA) or Doric (MFP_200/230/900-0.22_2m_FCM-MF). The subjects were on a treadmill consisting of a disk with a rotary encoder that pulsed 24 times per rotation. The mice were placed at about 7 cm from the center of the wheel. Thus, the distance walked with each tick was approximately 1.83 cm. As this could increase variability of the estimated run start time, we averaged 11-28 running bouts from each animal. Sensory stimuli were delivered as in two photon imaging experiments.

### Calcium imaging analysis

Initial image processing including correcting motion artifacts and drift in the imaging plane were done using the TurboReg plug-in in ImageJ (NIH) for 2-photon data or Mosaic (Inscopix) for miniature endoscope data. Imaging data were down-sampled spatially by 2*2 (2-photon) or 3*3 (miniature endoscope) binning to reduce file size. Cell outlines were drawn manually, and, in head-fixed 3-D 5-plane imaging, outlines containing the same cell were tracked across planes. Mean intensity within each region of interest (ROI) was used to generate *F_raw_*. The local neuropil signal was estimated from a neuropil mask containing the third to the sixth nearest pixels outside the outline that did not contain other ROIs, which was used to generate a mean neuropil signal *F_np_*. ROI specific signals were calculated as *F_raw_* –*F_np_*. For 2-photon imaging the ROI signals in neighbouring planes that clearly originated from the same cell were averaged to get the cell specific signal. The ROI specific signal was corrected for photobleaching in miniature endoscope data by computing ΔF/F_0_ using a median over a 100 s moving window as F_0_, to get the cell specific signal. Cell specific signals were Z-scored for plotting and analysis.

To define cell types, noise was reduced by smoothing *F_raw_* and *F_np_* with a 3 frame moving average, then average activity for each cell specific signal was calculated from bouts aligned to the onset of locomotion (Figure 2C). Baseline and 2 s.d. threshold were calculated from 10 – 6 s before locomotion onset. Extrema were found during the period 6 s before to 16 s after locomotion onset, and minimum and maximum responses were computed as 1 s averages around the extrema. If a cell had only a maximum above the 2 s.d. threshold from baseline it was called an ON cell, if it only had a minimum below 2 s.d. from baseline it was an OFF cell, if both of the previous conditions were true the cell was either an up-down or down-up cell (depending on the sequence of the extrema), and otherwise it was called not modulated. Onset times relative to movement onset (Figure 2P,Q) were calculated as the point where cell specific signal deviated at least 1 s.d. and stayed for at least 2 s at least 1 s.d. deviated from baseline. To calculate fractions in Figure 2R,S, we used a cutoff of mean + 2 s.d. from the whole recording for each cell to define active frames. Also, as locomotion-related activity preceded and followed locomotion epochs by some seconds, we defined movement frames for this analysis as those during movement and 3 s before and after.

In Figure 2T ON and up-down cells with the longest onset lead time were selected from previous imaging data and their average activity in each frame and the two preceding frames is plotted. In Figure 6E ON cell activity was similarly averaged across each frame and the preceding two frames.

Fiber photometry data in Figure 5J-L was corrected for photobleaching by computing ΔF/F_0_ using a mean over a 2 s period before each stimulus as F_0_. Fiber photometry data in Figure 5M was corrected for photobleaching by computing ΔF/F_0_ using a mean over a 240 s moving window as F_0_. ΔF/F_0_ was either averaged across sensory trials and smoothed with a 5 sample sliding average (Figure 5J-L), or averaged across locomotion onsets and smoothed with a 50 sample sliding average (Figure 5M). Signal onset was defined as first frame where signal increased 1 s.d. above baseline (10-6 s before locomotion, or 2.5-0 s before sensory stimulus) for at least 0.2 s.

### Optogenetics and O-DTR ablation experiments

Optogenetic manipulations were performed in the same head-fixed apparatus that was used for 2-photon imaging. Two 532 nm green lasers (Laserglow) were used to deliver light through shielded 0.2 mm diameter optic fibers to the bilateral implants. Light intensity out of the fiber was set to 3 mW for ArchT stimulation (constant on for 10, 12.05 or 30 s as indicated) and 20 mW for C1V1 stimulation (10 ms pulses for 2.5 s at indicated frequency). Orexin neurons in O-DTR mice were deleted by two injections of 150 ng diphtheria toxin (Sigma D0564) diluted to 1μg/ml in saline, administered i.p. on day 0 and day 2. Recordings for DT+ conditions were started on day 12. WT control mice were treated identically in parallel. In experiments in Figure 4 and 6F-O, the trial would not start until the mouse was stationary for at least 5 s. In addition, trials were spaced by 15 s + randomized 0.1-10 s intervals (Figure 4) or by 25 s + randomized 0.1-25 s intervals (Figure 6F-O) and the order of laser-on and laser-off (catch) trials was randomized in each block (consisting of one of each trial). In Figure 6F-J laser was turned on 50 ms before stimulus presentation. In order to limit the total duration of each recording session, due to the number of stimuli and length of intervals, the datasets for ArchT and O-DTR experiments were combined across two recording sessions from each mouse. As orexin neuron deletion can lead to a cataplectic phenotype, we monitored video footage of the mice during locomotion recordings to check for signs of abrupt sleep attacks or loss of muscle tone. However, no atonia was seen throughout recordings in the O-DTR DT+ and ArchT laser+ conditions, and in the ArchT experiments, there was no effect on speed or duration of movement bouts, suggesting strongly that cataplectic attacks did not occur in these recordings. This is also consistent with the rarity of cataplectic attacks in the absence emotional triggers (Hara et al., 2001; Scammell et al., 2009), and a previously demonstrated lack of effect of transient orexin-ArchT inhibition during the dark phase (Tsunematsu et al., 2013). Care was taken to record each mouse at the same time of day to minimize circadian effects. Before every experiment mice were allowed to habituate to head-fixation and stimulus presentation and or optical manipulation for 5-10 min.

### Preparation of acute slices

Coronal brain slices from P60-180 animals were prepared after instant cervical dislocation and decapitation. The brain was rapidly dissected and cooled in continuously gassed (95% O_2_ and 5% CO_2_), icy cutting solution containing (in mM): 90 N-methyl-D-glucamine, 20 HEPES, 110 HCl, 3 KCl, 10 MgCl_2_, 0.5 CaCl_2_, 1.1 NaH_2_PO_4_, 25 NaHCO_3_, 3 pyruvic acid, 10 ascorbic acid and 25 D-glucose. 350 μm thick coronal brain slices were cut on a vibratome (Campden) and allowed to recover for 5-15 min at 37 °C in cutting solution, followed by 45-55 min at 22 °C in artificial cerebrospinal fluid (ACSF) containing (in mM): 126 NaCl, 3 KCl, 2 MgSO_4_, 2 CaCl_2_, 1.1 NaH_2_PO_4_, 26 NaHCO_3_, 0.1 pyruvic acid, 0.5 L-glutamine, 0.4 ascorbic acid and 25 D-glucose, continuously gassed with 95% O_2_ and 5% CO_2_.

### Slice electrophysiology

Patch clamp recordings were performed in a submerged chamber with 3-5 ml/min superfusion with ACSF, continuously gassed with 95% O_2_ and 5% CO_2_. 3-6 MOhm patch pipettes were filled with intracellular solution containing (in mM): 130 K-gluconate, 5 NaCl, 2 MgSO_2_, 10 HEPES, 0.1 EGTA, 4 Mg-ATP, 0.4 Na-GTP, 2 pyruvic acid, 0.1 Alexa-594, 0.1% biocytin, and ~10 mM KOH (to set pH to 7.3). Whole cell recordings were not analysed if the access resistance was above 25 MOhm. Voltage recordings were sampled at 10 kHz and low-pass filtered at 3 kHz with HEKA EPC10 usb amplifiers and acquired with HEKA patchmaster software. ArchT and C1V1 were stimulated with green light from a xenon lamp (Sutter lambda 4DG controlled from HEKA patchmaster) through a TRITC-filter. Concurrent GCaMP6s fluorescence recordings were performed using the miniature endoscope with a GRIN lens implant recovered from a successful in vivo recording. This assembly was mounted on a micromanipulator and moved into focus at about 350 μm away from the patch clamped cell. Patch clamp data were analyzed in Matlab.

### Immunohistochemistry

Mice were overdosed with ketamine/xylazine (100mg/kg and 10mg/kg respectively) and transcardially perfused with 4% PFA. Brains were post-fixed for 24 h. Coronal brain slices were sectioned at 50 μm using a cryostat or a vibratome. Sections were blocked in PBS with 0.3% Triton X-100 and 1% bovine serum albumin (blocking solution) for 1 h, incubated with goat anti-orexin (1:1000; Santa Cruz) over-night, washed, incubated with Alexa 647 conjugated donkey anti-goat (1:1000; Invitrogen) for 3.5 h, washed and mounted. Antibodies were applied blocking solution. Confocal micrographs were acquired on an Olympus FV1000 and merged in imageJ.

### Statistics

All data are shown as mean ± SEM unless stated otherwise. Statistical significance was determined by paired Student t-test or Wilcoxon rank sum test as stated. All statistics were performed using statistical functions in Matlab. Sigmoid fits in Figure 3C and E are based on modified Hill equations 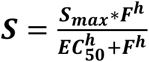 (Figure 3C) or 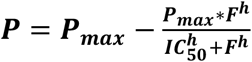 (Figure 3E), where S is spike output, F is light pulse frequency, P is movement probability, EC_50_ is half-maximal response and IC_50_ half-maximal decrease. Fitting was done using custom routines in Matlab.

---

**Supplementary figure 1.**
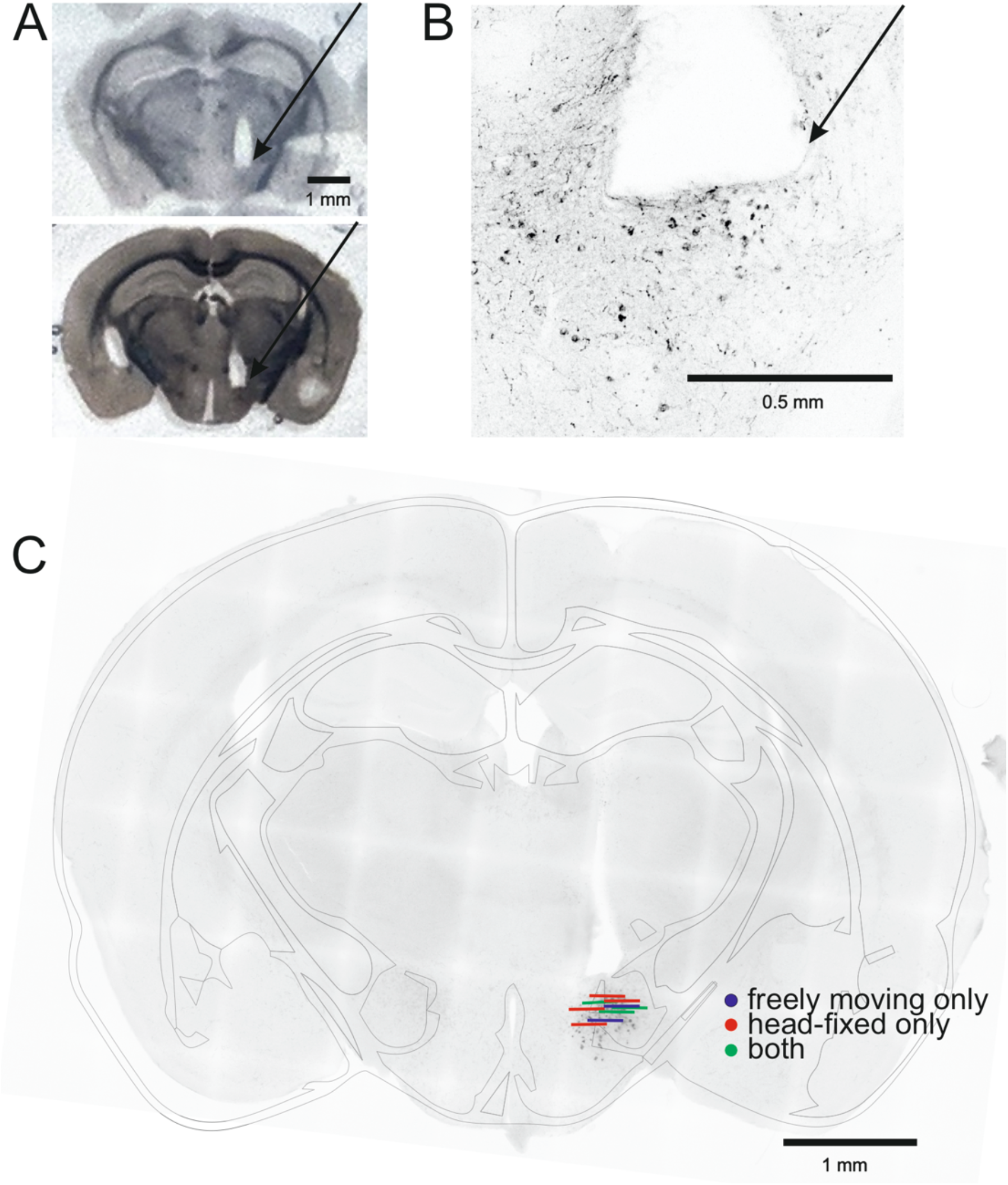
GRIN lens placement. A, Coronal sections from two mice illustrating GRIN lens placement in LH (arrow). B, Confocal micrograph of orexin cells and GRIN lens track (arrow) in coronal section. C, Reconstructed approximate imaging plane locations in the 9 mice used in this study. Blue = 2 mice that were recorded only with miniature endoscope in the freely-moving paradigm, red = 4 mice that were recorded only with 2-photon microscopy in head-fixed paradigms, green = 3 mice that were recorded in both paradigms.

## Acknowledgements

We thank Lise T. Jensen and Lars Fugger for sharing O-DTR mice and Takeshi Sakurai for sharing the orexin promoter used for generating orexin neuron specific constructs. We thank Alan Ling and Adam Hurst at the Francis Crick Institute mechanical workshop for making recording arenas, staff at the Crick biological research facility for animal husbandry, staff at Vigene for technical assistance with viruses, and Bruno Pichler and Dale Elgar for microscope design. We thank Troy Margrie for sharing equipment and all Burdakov lab members for useful comments and help. This project has received funding from the European Union’s Horizon 2020 research and innovation programme under the Marie Skłodowska-Curie grant agreement DRIVOME (grant agreement 701986). This work was funded by The Francis Crick Institute, which receives its core funding from Cancer Research UK, the UK Medical Research Council.

## Author contributions

Conceptualization: MMK, DB, AA. Methodology: MMK, DB, CS, EFB, JAG, PV. Software: MMK, CS, EFB. Analysis: MMK. Investigation: MMK, CS, EFB, JAG. Resources: DB. Writing – original draft preparation: MMK, DB. Writing – review and editing: all authors. Funding acquisition: MMK, DB.

## References

Adamantidis, A.R., Zhang, F., Aravanis, A.M., Deisseroth, K., de Lecea, L., 2007. Neural substrates of awakening probed with optogenetic control of hypocretin neurons. Nature. 450, 420–4. https://doi.org/10.1038/nature06310

Allen, W.E., Kauvar, I.V., Chen, M.Z., Richman, E.B., Yang, S.J., Chan, K., Gradinaru, V., Deverman, B.E., Luo, L., Deisseroth, K., 2017. Global Representations of Goal-Directed Behavior in Distinct Cell Types of Mouse Neocortex. Neuron 94, 891–907.e6. https://doi.org/10.1016/j.neuron.2017.04.017

Apergis-Schoute, J., Iordanidou, P., Faure, C., Jego, S., Schöne, C., Aitta-Aho, T., Adamantidis, A., Burdakov, D., 2015. Optogenetic evidence for inhibitory signaling from orexin to MCH neurons via local microcircuits. J. Neurosci. Off. J. Soc. Neurosci. 35, 5435–41. https://doi.org/10.1523/JNEUROSCI.5269-14.2015

Arber, S., Costa, R.M., 2018. Connecting neuronal circuits for movement. Science 360, 1403–1404. https://doi.org/10.1126/science.aat5994

Arrigoni, E., Chee, M.J.S., Fuller, P.M., 2018. To eat or to sleep: That is a lateral hypothalamic question. Neuropharmacology. https://doi.org/10.1016/j.neuropharm.2018.11.017

Bonnavion, P., Jackson, A.C., Carter, M.E., de Lecea, L., 2015. Antagonistic interplay between hypocretin and leptin in the lateral hypothalamus regulates stress responses. Nat. Commun. 6, 6266. https://doi.org/10.1038/ncomms7266

Burton, M.J., Rolls, E.T., Mora, F., 1976. Effects of hunger on the responses of neurons in the lateral hypothalamus to the sight and taste of food. Exp. Neurol. 51, 668–677.

Chen, T.-W., Wardill, T.J., Sun, Y., Pulver, S.R., Renninger, S.L., Baohan, A., Schreiter, E.R., Kerr, R.A., Orger, M.B., Jayaraman, V., Looger, L.L., Svoboda, K., Kim, D.S., 2013. Ultrasensitive fluorescent proteins for imaging neuronal activity. Nature 499, 295–300. https://doi.org/10.1038/nature12354

Crochet, S., Lee, S.-H., Petersen, C.C.H., 2019. Neural Circuits for Goal-Directed Sensorimotor Transformations. Trends Neurosci. 42, 66–77. https://doi.org/10.1016/j.tins.2018.08.011

Cui, G., Jun, S.B., Jin, X., Pham, M.D., Vogel, S.S., Lovinger, D.M., Costa, R.M., 2013. Concurrent activation of striatal direct and indirect pathways during action initiation. Nature 494, 238–242. https://doi.org/10.1038/nature11846

da Silva, J.A., Tecuapetla, F., Paixão, V., Costa, R.M., 2018. Dopamine neuron activity before action initiation gates and invigorates future movements. Nature 554, 244–248. https://doi.org/10.1038/nature25457

Ferezou, I., Haiss, F., Gentet, L.J., Aronoff, R., Weber, B., Petersen, C.C.H., 2007. Spatiotemporal dynamics of cortical sensorimotor integration in behaving mice. Neuron 56, 907–923. https://doi.org/10.1016/j.neuron.2007.10.007

González, J.A., Iordanidou, P., Strom, M., Adamantidis, A., Burdakov, D., 2016a. Awake dynamics and brain-wide direct inputs of hypothalamic MCH and orexin networks. Nat. Commun. 7, 11395. https://doi.org/10.1038/ncomms11395

González, J.A., Jensen, L.T., Iordanidou, P., Strom, M., Fugger, L., Burdakov, D., 2016b. Inhibitory Interplay between Orexin Neurons and Eating. Curr. Biol. 26, 2486–2491. https://doi.org/10.1016/j.cub.2016.07.013

Hara, J., Beuckmann, C.T., Nambu, T., Willie, J.T., Chemelli, R.M., Sinton, C.M., Sugiyama, F., Yagami, K., Goto, K., Yanagisawa, M., Sakurai, T., 2001. Genetic ablation of orexin neurons in mice results in narcolepsy, hypophagia, and obesity. Neuron 30, 345–354.

Hassani, O.K., Krause, M.R., Mainville, L., Cordova, C.A., Jones, B.E., 2016. Orexin Neurons Respond Differentially to Auditory Cues Associated with Appetitive versus Aversive Outcomes. J. Neurosci. 36, 1747–1757. https://doi.org/10.1523/JNEUROSCI.3903-15.2016

Hu, B., Yang, N., Qiao, Q.-C., Hu, Z.-A., Zhang, J., 2015. Roles of the orexin system in central motor control. Neurosci. Biobehav. Rev. 49, 43–54. https://doi.org/10.1016/j.neubiorev.2014.12.005

Inutsuka, A., Yamashita, A., Chowdhury, S., Nakai, J., Ohkura, M., Taguchi, T., Yamanaka, A., 2016. The integrative role of orexin/hypocretin neurons in nociceptive perception and analgesic regulation. Sci. Rep. 6, 29480. https://doi.org/10.1038/srep29480

Iyer, M., Essner, R.A., Klingenberg, B., Carter, M.E., 2018. Identification of discrete, intermingled hypocretin neuronal populations. J. Comp. Neurol. 526, 2937–2954. https://doi.org/10.1002/cne.24490

Jordan, L.M., 1998. Initiation of locomotion in mammals. Ann. N. Y. Acad. Sci. 860, 83–93.

Kerman, I.A., Bernard, R., Rosenthal, D., Beals, J., Akil, H., Watson, S.J., 2007. Distinct populations of presympathetic-premotor neurons express orexin or melanin-concentrating hormone in the rat lateral hypothalamus. J. Comp. Neurol. 505, 586–601. https://doi.org/10.1002/cne.21511

Kosse, C., Schöne, C., Bracey, E., Burdakov, D., 2017. Orexin-driven GAD65 network of the lateral hypothalamus sets physical activity in mice. Proc. Natl. Acad. Sci. 114, 4525–4530. https://doi.org/10.1073/pnas.1619700114

Lee, M.G., Hassani, O.K., Jones, B.E., 2005. Discharge of Identified Orexin/Hypocretin Neurons across the Sleep-Waking Cycle. J. Neurosci. 25, 6716–6720. https://doi.org/10.1523/JNEUROSCI.1887-05.2005

Li, Y., Zeng, J., Zhang, J., Yue, C., Zhong, W., Liu, Z., Feng, Q., Luo, M., 2018. Hypothalamic Circuits for Predation and Evasion. Neuron 97, 911–924.e5. https://doi.org/10.1016/j.neuron.2018.01.005

Mahler, S.V., Moorman, D.E., Smith, R.J., James, M.H., Aston-Jones, G., 2014. Motivational activation: a unifying hypothesis of orexin/hypocretin function. Nat. Neurosci. 17, 1298–303. https://doi.org/10.1038/nn.3810

Mickelsen, L.E., Bolisetty, M., Chimileski, B.R., Fujita, A., Beltrami, E.J., Costanzo, J.T., Naparstek, J.R., Robson, P., Jackson, A.C., 2019. Single-cell transcriptomic analysis of the lateral hypothalamic area reveals molecularly distinct populations of inhibitory and excitatory neurons. Nat. Neurosci. 22, 642–656. https://doi.org/10.1038/s41593-019-0349-8

Mickelsen, L.E., Kolling, F.W., Chimileski, B.R., Fujita, A., Norris, C., Chen, K., Nelson, C.E., Jackson, A.C., 2017. Neurochemical Heterogeneity Among Lateral Hypothalamic Hypocretin/Orexin and Melanin-Concentrating Hormone Neurons Identified Through Single-Cell Gene Expression Analysis. eneuro 4, ENEURO.0013-17.2017. https://doi.org/10.1523/ENEURO.0013-17.2017

Mileykovskiy, B.Y., Kiyashchenko, L.I., Siegel, J.M., 2005. Behavioral Correlates of Activity in Identified Hypocretin/Orexin Neurons. Neuron 46, 787–798. https://doi.org/10.1016/j.neuron.2005.04.035

Mohajerani, M.H., Chan, A.W., Mohsenvand, M., LeDue, J., Liu, R., McVea, D.A., Boyd, J.D., Wang, Y.T., Reimers, M., Murphy, T.H., 2013. Spontaneous cortical activity alternates between motifs defined by regional axonal projections. Nat. Neurosci. 16, 1426–1435. https://doi.org/10.1038/nn.3499

Mora, F., Rolls, E.T., Burton, M.J., 1976. Modulation during learning of the responses of neurons in the lateral hypothalamus to the sight of food. Exp. Neurol. 53, 508–519.

Moser, E.I., Moser, M.-B., McNaughton, B.L., 2017. Spatial representation in the hippocampal formation: a history. Nat. Neurosci. 20, 1448–1464. https://doi.org/10.1038/nn.4653

Rolls, E.T., Burton, M.J., Mora, F., 1976. Hypothalamic neuronal responses associated with the sight of food. Brain Res. 111, 53–66.

Saper, C.B., Scammell, T.E., Lu, J., 2005. Hypothalamic regulation of sleep and circadian rhythms. Nature 437, 1257–1263. https://doi.org/10.1038/nature04284

Scammell, T.E., Willie, J.T., Guilleminault, C., Siegel, J.M., 2009. A Consensus Definition of Cataplexy in Mouse Models of Narcolepsy 32, 6.

Schöne, C., Cao, Z.F.H., Apergis-Schoute, J., Adamantidis, A., Sakurai, T., Burdakov, D., 2012. Optogenetic probing of fast glutamatergic transmission from hypocretin/orexin to histamine neurons in situ. J. Neurosci. Off. J. Soc. Neurosci. 32, 12437–43. https://doi.org/10.1523/JNEUROSCI.0706-12.2012

Schöne, C., Venner, A., Knowles, D., Karnani, M.M., Burdakov, D., 2011. Dichotomous cellular properties of mouse orexin/hypocretin neurons. J. Physiol. 589, 2767–79. https://doi.org/10.1113/jphysiol.2011.208637

Sinnamon, H.M., 1993. Preoptic and hypothalamic neurons and the initiation of locomotion in the anesthetized rat. Prog. Neurobiol. 41, 323–344.

Svoboda, K., Li, N., 2018. Neural mechanisms of movement planning: motor cortex and beyond. Curr. Opin. Neurobiol. 49, 33–41. https://doi.org/10.1016/j.conb.2017.10.023

Takahashi, K., Lin, J.-S., Sakai, K., 2008. Neuronal activity of orexin and non-orexin waking-active neurons during wake-sleep states in the mouse. Neuroscience 153, 860–870. https://doi.org/10.1016/j.neuroscience.2008.02.058

Tsunematsu, T., Tabuchi, S., Tanaka, K.F., Boyden, E.S., Tominaga, M., Yamanaka, A., 2013. Long-lasting silencing of orexin/hypocretin neurons using archaerhodopsin induces slow-wave sleep in mice. Behav. Brain Res. 255, 64–74. https://doi.org/10.1016/j.bbr.2013.05.021

Tyree, S.M., Borniger, J.C., de Lecea, L., 2018. Hypocretin as a Hub for Arousal and Motivation. Front. Neurol. 9, 413. https://doi.org/10.3389/fneur.2018.00413

Vélez-Fort, M., Bracey, E.F., Keshavarzi, S., Rousseau, C.V., Cossell, L., Lenzi, S.C., Strom, M., Margrie, T.W., 2018. A Circuit for Integration of Head-and Visual-Motion Signals in Layer 6 of Mouse Primary Visual Cortex. Neuron 98, 179–191.e6. https://doi.org/10.1016/j.neuron.2018.02.023

Yamanaka, A., Beuckmann, C.T., Willie, J.T., Hara, J., Tsujino, N., Mieda, M., Tominaga, M., Yagami, K. ichi, Sugiyama, F., Goto, K., Yanagisawa, M., Sakurai, T., 2003. Hypothalamic orexin neurons regulate arousal according to energy balance in mice. Neuron 38, 701–713.

